# Basecalling for DNA Storage

**DOI:** 10.1101/2025.07.27.667078

**Authors:** Advait Menon, Samira Brunmayr, Omer Sella, Thomas Heinis

## Abstract

DNA is a promising medium for data storage with its high information density and stability. To retrieve information stored in DNA, sequencing technologies are used to read the encoded bases. The raw signals from sequencing are mapped to a sequence of {A,C,T,G} by machine learning algorithms known as basecallers. Currently, basecallers are optimised mainly on biological DNA instead of focusing on characteristics or constraints unique to data-encoding DNA. Taking advantage of these unique artificial features to fine-tune and adapt the architecture, we demonstrate for the first time that DNA coding scheme constraints can be leveraged to optimise basecallers. Using low-rank adaptation on the basecalling model, we achieve substantial gains with high resource efficiency. Additionally, constraint-aware beam search provides improvements without requiring model retraining.

## Introduction

The global demand for data storage is increasing exponentially, with worldwide data creation expected to exceed 394 zettabytes by 2028 [1]. Current storage technologies such as magnetic tapes and optical disks are nearing their information density limits [2], prompting the exploration of alternative, long-term storage media. DNA has emerged as a promising candidate for archival data storage due to its extremely high density, with a theoretical maximum of 455 exabytes per gram of single-stranded DNA [3], and its stability over thousands of years under the right conditions, with little-to-no maintenance [4].

At present, a limitation of the DNA storage pipeline lies in the sequencing step, which is the process of reading the DNA [5]. One of the most promising readout methods for DNA storage systems is nanopore sequencing, where a DNA molecule is pulled through a nanopore, generating an electrical signal. This signal is modulated by the chemical composition of each base as it passes through the nanopore, and is then translated into a textual sequence of nucleotides {A,C,T,G} by algorithms known as basecallers [6]. While it is faster, cheaper and more portable than its alternatives [7], nanopore sequencing suffers from a drastically lower read accuracy (87-98%, whereas first- and second-gen sequencing methods can achieve >99.9%, and third-gen achieves >99%) [8].

Although redundancy and error-correcting codes can partially mitigate thisissue [9, 10], such approaches are mostly effective with first- and second-generation sequencing technologies, due to their low and substitution-dominated errors [11, 12]. The significantly lower read accuracy of nanopore sequencing often exceeds their error-handling capabilities, rendering them less effective. Decoding error-correcting codes can also be computationally intensive, particularly in high-error environments (like those encountered with nanopore sequencing) [13]. Furthermore, error-correction and redundancy mechanisms often trade robustness, for reduced information density.

These challenges highlight the need to improve the accuracy of basecalling itself, rather than relying solely on downstream correction. Modern basecalling tools such as SACall [14], Bonito [15], and Dorado [16] use deep neural networks trained on biological DNA. As a result, they may perform suboptimally when applied to synthetic, data-encoding DNA. Basecallers generally perform best when trained on species-specific data as shown by Pagès-Gallego et al. [17] and Ferguson et al. [5]. Our work considers data-encoding DNA as its own “species”, with distinct statistical properties that current models are not optimised for.

In this paper, we evaluate whether incorporating constraints unique to DNA coding schemes, such as limiting the number of repeated bases and fixing the length of sequences, can improve basecalling accuracy for data-carrying DNA. We investigate two strategies: (1) fine-tuning a pre-trained basecaller (SACall) using synthetic nanopore signals from coded DNA sequences, including efficient fine-tuning via low-rank adaptation (LoRA); and (2) modifying the decoding process with a constraint-aware beam search algorithm. Together, these approaches aim to improve accuracy and data recovery without increasing computational overhead or relying on additional hardware, as opposed to state of the art basecalling practice [18]. Our results show that significant performance gains can be achieved by aligning basecalling methods with the structure of data-carrying DNA, increasing read accuracy from 85.8% to over 95% through fine-tuning, and improving it by 1.7% without requiring any retraining by using constraint-aware decoding.

## Materials and Methods

### The benchmark

SACall was chosen as the benchmark basecaller in this work. We chose to reimplement and train this model from scratch with PyTorch, re-using some modules from the original source code, to maintain full control over the training process. The reason why we chose SACall is due to its state-of-the-art performance, with a match rate second only to Bonito [17] and, unlike Bonito, its publicly available training data - making it a better candidate, especially when checking implementation correctness. Fig 1 outlines the model architecture. The encoder comprises of a series of convolutional layers followed by transformer encoder layers. The decoder consists of a single linear layer. During training, we employed CTC loss to enable the model to learn alignments and map variable-length nanopore signals to DNA sequences without requiring explicit alignment. For inference, the output tensor is decoded using beam search (from the fast-ctc-decode library [19]) with a beam width of 3, balancing accuracy and computation speed.

**Fig 1.**
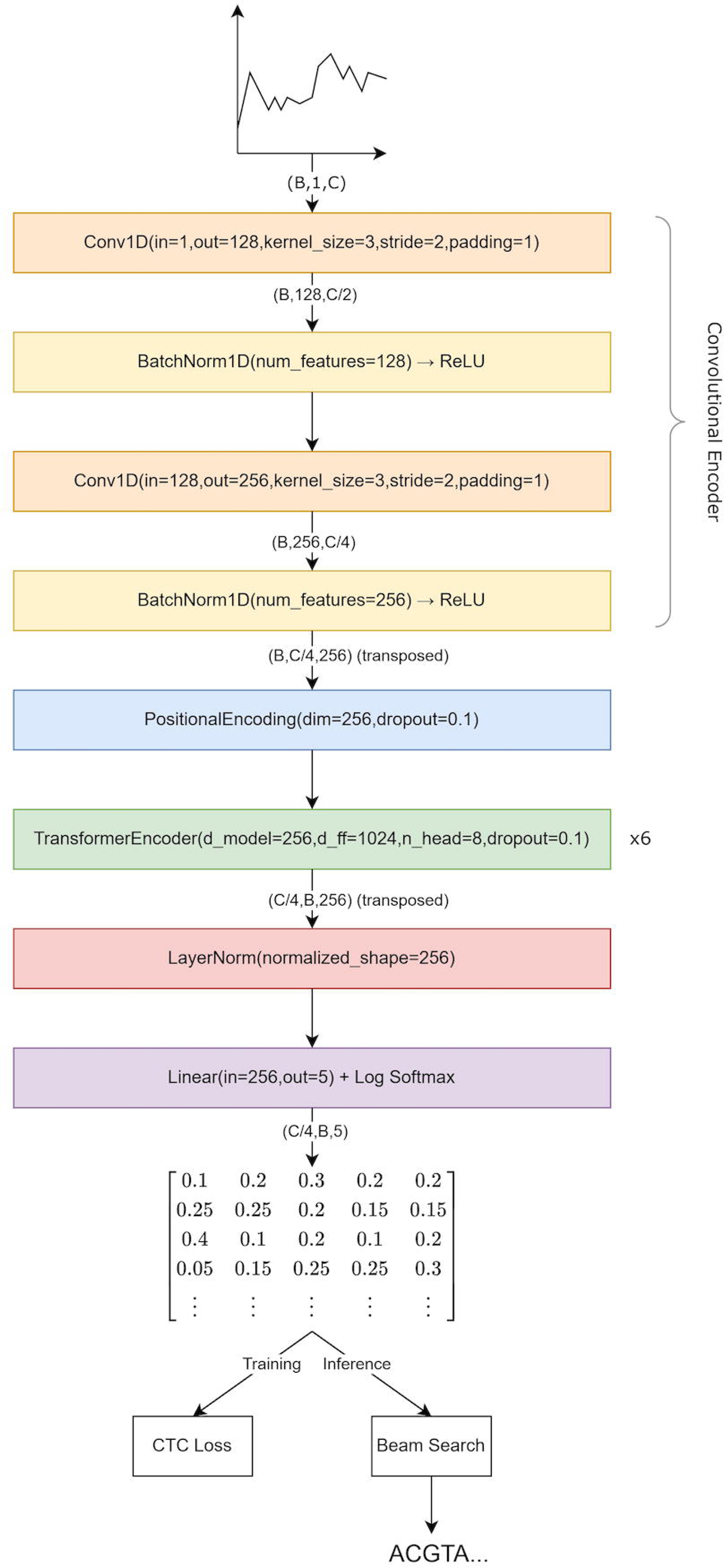
Architecture of the benchmark basecaller, based on SACall [14]. *B* denotes the batch size and *C* denotes the chunk size.

### Data Pre-processing and Pre-Training

The benchmark model was pre-trained on the same bacterial species dataset as the original SACall model, provided by Wick et al. [18]. To pre-process the dataset, the approach specified in Pagès-Gallego et al. [17] was used, adapting their framework to implement and pre-train the benchmark model. First, all nanopore reads were annotated with their reference sequences using Tombo [20]. Next, Tombo’s resquiggle algorithm was executed to define a new assignment from squiggles to reference sequences. This ensures that sequencing errors in the reference sequences were mitigated and improved the overall quality of the training data. Furthermore, to minimise the effect of signal variations across batches, the signals were normalised with median shift and median absolute deviation scale parameters [14]:

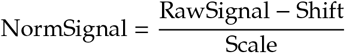

Finally, the annotated raw signals were processed into non-overlapping, fixed-size chunks of 2000 signal values, to reduce memory consumption and ensure consistent sequence lengths [21], and divided into training and validation sets with an 80/20 split.

The model was trained with an Adam optimiser [22] with an initial learning rate of and a gradual warm-up scheduler [23] with 5000 warm-up steps, followed by a cosine annealing scheduler. The benchmark model was trained for 4 epochs on a single Tesla A30 24GB (CUDA 12.4) for 56 hours with a batch size of 64. For reproducibility, we set all seeds before training.

### Fine-Tuning

The extent to which the constraints on data-encoding DNA can be leveraged through fine-tuning of the benchmark model to improve accuracy was explored. Three fine-tuning methods were implemented:

1. **Whole-Model Fine-Tuning**: This fine-tunes all parameters of the pre-trained model on the simulated data-encoding DNA dataset. Although computationally expensive, it allows the entire network to adjust, thus serving as an upper bound in terms of model adaptability.
2. **Decoder Fine-Tuning**: This fine-tunes only the parameters in the final linear layer of the model and freezes all others (Fig 2B). The goal is to assess whether adapting only the final projection to the output logits is sufficient to capture the characteristics of data-encoding DNA.
3. **Low-Rank Adaptations (LoRA)**: Low-Rank Adaptations (LoRA), introduced by Hu et al. [24], is a form of parameter-efficient fine-tuning (PEFT) [25] that allows efficient adaptation, while maintaining the priors learned during pre-training on bacterial DNA (Fig 2C). As far as we are aware, no prior work has used LoRA for basecalling, making this a novel application.

**Fig 2.**
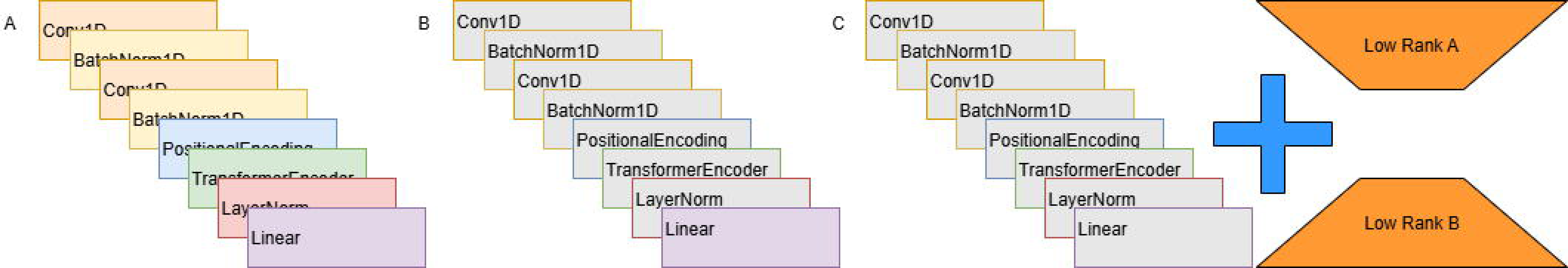
Fine tuning approaches in this work. Weights that are kept frozen are shown in Grey. (A) Whole model approach. All model weights are subject to fine tuning. (B) Last layer tuning. The weights of the remaining layers are kept frozen. (C) Low rank adaptation. Model weights are frozen, and a low rank matrtix is fine tuned and added to the model.

Each model was fine-tuned for 45,000 epochs, with a batch size of 64 on a single Tesla A30 24GB (CUDA 12.4). The pre-trained SACall model was used as the baseline for fine-tuning and multiple LoRA models were fine-tuned, varying the rank.

Currently, there are no publicly available well-curated datasets of data-encoding DNA nanopore reads. We hence encoded data from the CIFAR-10 dataset [26] using Goldman coding [9], and then mapped the encoded data to simulated raw nanopore squiggles using Squigulator [27]. We chose to use Goldman coding since, by construction, it imposes three key constraints which are needed to ensure stable DNA structures [28]:

- **Homopolymer constraint**, which limits the maximum length of repeated bases (e.g., AAAA)
- **Length constraint**, which fixes the sequences length to 117 nucleotides
- **Base composition constraint**, which ensures an approximately uniform distribution over the four nucleotides {A, C, T, G}.

In order to maintain consistency with the original bacterial data used for pre-training, the simulated data was pre-processed in a similar manner, i.e., by annotating and re-squiggling with Tombo, normalising the signals, and splitting the raw signals into non-overlapping chunks of 2,000 signal values.

### Beam Search Modifications

A constraint-aware decoder was designed, making use of known characteristics of the Goldman code in the beam search algorithm. This constraint-aware beam search was implemented based on a fork of Oxford Nanopore Technologies’ fast-ctc-decode [19] library, which is commonplace among standard CTC basecallers due to its high speed.

#### Homopolymer constraint

The homopolymer constraint on Goldman-encoded data enforces a maximum homopolymer length of 1 on generated DNA sequences.

The default beam search scores the various hypothesis extensions by descending order of probability. We defined a new score that subtracts a constant penalty from the probability of any extension exceeding a provided maximum homopolymer length:

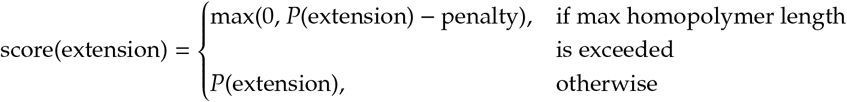

#### Length constraint

Goldman coding constructs DNA sequences that are 117 nucleotides long.

We explored two methods for leveraging the length constraint, in order to guide the beam search towards nucleotide sequences of length closer to the desired 117nt.

We defined a new scoring, whereby extensions are penalised by their normalised absolute difference to the desired length. This ensures that extensions whose length differs greatly from the desired length are penalised more.

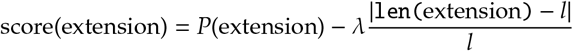

where λ≥0 is a scaling coefficient and *l* = 117nt is the desired length of the sequence.

Our second approach involved discarding any extensions which stray too far from the desired length at a given time step. We compute the length per number of time steps for the desired sequence length:

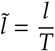

where *l* = 117 is the desired length, and T is the total number of time steps in the model output tensor. At each time step in the beam search, we compute the length per time step of the extension we are considering and then discard the extension if:

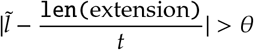

where *t* is the time step/beam search iteration and θ is a provided threshold.

#### Base distribution constraint

Assuming an independent and uniform distribution over trits at each index in the base-3 encoded data, the distribution of the four bases in the encoded payload is approximately uniform. Although the distribution of trits in a given file is unknown, we hypothesise that guiding the distribution of basecalled sequences towards the uniform distribution will yield alignment rate and accuracy improvements over the baseline beam search algorithm.

We use the Kullback-Leibler (KL) divergence to quantify how close the distribution of bases in a nucleotide sequence is to the uniform distribution:

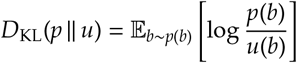

where

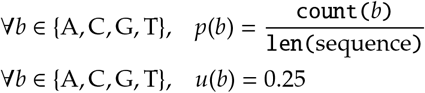

We incorporated the KL-divergence into the extension scoring function:

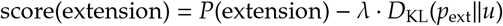

where λ > 0 is a scaling coefficient, *p*_ext_ is the base distribution of the extension and *u* is the uniform distribution.

We also explored discarding any hypothesis extensions that would result in an imbalanced distribution of bases. More formally, at each beam search time step/iteration, rather than considering all bases for extending the current sequence, our modified algorithm only considers extensions for bases that are under-represented. A base *b* is under-represented if:

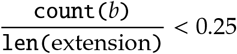

Consider the predicted base distribution at time step *t*:

*p*(A) = 0.8, *p*(C) = 0.1, *p*(G) = 0.05, *p*(T) = 0.05. An issue with this strategy is that if ‘A’ would be over-represented in the proposed extension, we discard it completely, despite the model output distribution heavily favouring it. To address this, we took inspiration from Sneddon et al. [21] and only discard the extension if both: a) the KL divergence to the uniform is greater than a threshold θ and b) the Shannon entropy (quantifying the uncertainty) is above a threshold τ.

The Shannon entropy of a distribution *p*(*x*) for random variable *X* is:

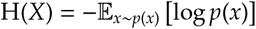

We considered an extension to be uncertain if its entropy was greater than 80% of the upper-bound, in other words we set the threshold τ = 0.8 · log 4.

### Evaluation metrics

To evaluate the model, we first set up our test set by encoding the same set of files used by Goldman et al. [9]. The details of the test set are shown in S1 Table. All reads within the test set are then basecalled, and the predicted sequences are aligned with their corresponding reference sequences using Mappy (the python bindings for Minimap2 [29]). This yields the number of matches (M), mismatches/substitutions (X), insertions (I) and deletions (D).

A match occurs when a predicted base aligns to the same base at the same position in the reference sequence. A mismatch (or substitution) occurs when the aligning bases are different. An insertion is an extra base present in the predicted sequence that does not align to a base in the reference sequence. A deletion is a base in the reference sequence that does not align to a base in the predicted sequence.

Provided that Mappy can successfully align the predicted sequence to the reference, we can define the following evaluation metrics:

#### Read Accuracy

Read accuracy is the proportion of matching bases with respect to the length of the reference sequence:

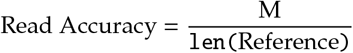

#### M*/*X*/*I*/*D Rates

Match/Mismatch/Insertion/Deletion rates are all computed with respect to the length of the alignment:

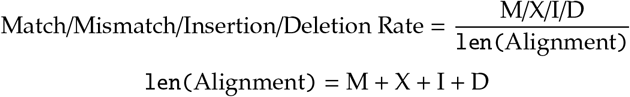

len(Alignment) = M + X + I + D

#### Alignment Rate

Alignment rate is the proportion of predicted sequences that were successfully aligned by Mappy:

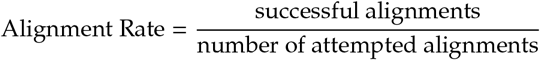

It is essential to report this metric alongside read accuracy, since a high read accuracy is meaningless if only a small portion of the total number of base calls could be aligned.

#### Payload Accuracy

While the metrics such as median read accuracy and alignment rate provide insight into basecaller performance on individual raw data signals, they do not quantify our ability to recover the entire encoded data. In Goldman coding, illustrated in Fig 3, the data is split into overlapping fragments of length 100nt (nucleotides) with a step size of 25nt. These fragments are sandwiched between two indexing information to enable data reconstruction. Although Goldman coding implements four-fold redundancy for the payload, there is no such mechanism in place for the strand indexing information. To address this, we assume that the strand index of each predicted DNA sequence is known. In other words, we disregard any erroneous bases in the indexing information of a sequence. We also assume knowledge of which portion of the predicted sequence aligns to the reference sequence. We then reconstruct the payload, using majority voting on each overlapping 25nt segment (breaking ties randomly).

**Fig 3.**
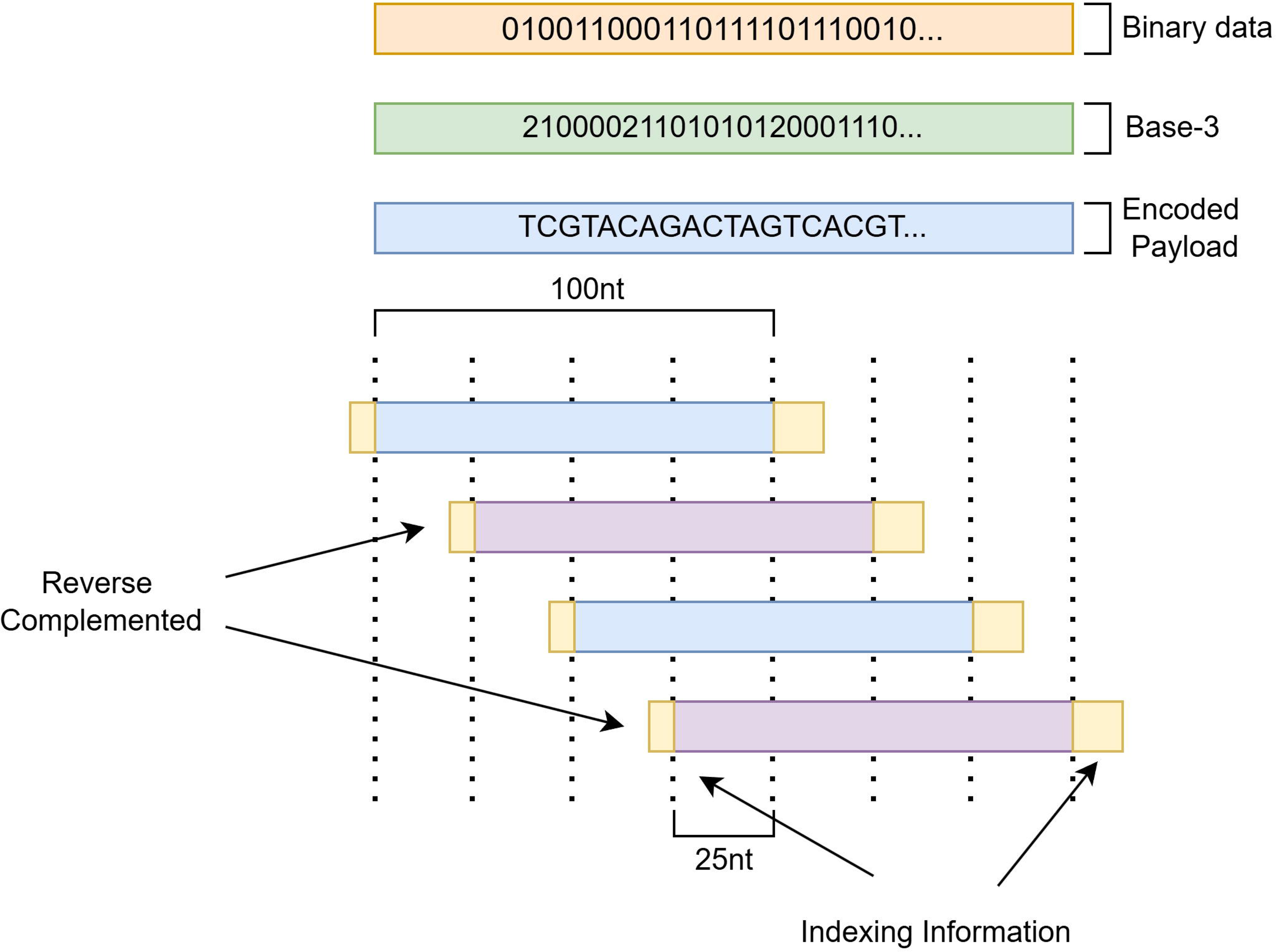
Goldman coding process, from binary data to the final DNA fragments (olignonucleotides). Each DNA fragment has a payload of 100nt (nucleotides) and 17nt of indexing information. Reproduced from [9].

We then define the *payload accuracy*:

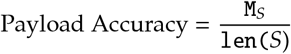

where M_*S*_ is the number of matching bases between the reference encoded payload *S* and a reconstruction of the encoded payload Ŝ from the basecalled nucleotide sequences.

## Results and Discussion

### Fine-Tuning Evaluation

Table 1 summarises the performance of each fine-tuning strategy on the synthetic evaluation dataset (the same set of files used by Goldman et al. [9]). It also shows the homopolymer, length and base distribution measurements to quantify how closely the constraints imposed by Goldman coding were adhered to by sequences that succeeded and failed (F) alignment. Table 2 displays the number of trainable parameters for each method.

As the experiment shows, all fine-tuned models outperformed the baseline, both in terms of performance metrics, and how well they adhered to the constraints.

Decoder fine-tuning performed the worst among the strategies while still outperforming the baseline by exceeding the alignment rate for almost all files by approximately 20%, median read accuracy by approximately 5% and payload accuracy by up to 13%. This demonstrates that despite having the fewest trainable parameters of the strategies employed, the decoder plays a meaningful role in basecaller prediction.

Both whole-model fine-tuning and LoRA strategies performed exceedingly well, achieving read accuracies >95%, alignment rate >94% and payload accuracy >94% (for ranks ≥ 2). Surprisingly, LoRA with rank ≥ 4 achieved greater payload accuracy than whole-model fine-tuning, despite having significantly fewer trainable parameters. In addition, the training curves for whole-model fine-tuning (S3 Fig) indicate that the model did converge. This suggests that fine-tuning the entire model can disrupt previously learned features — explaining its marginally worse performance compared to LoRA.

**Table 1.**
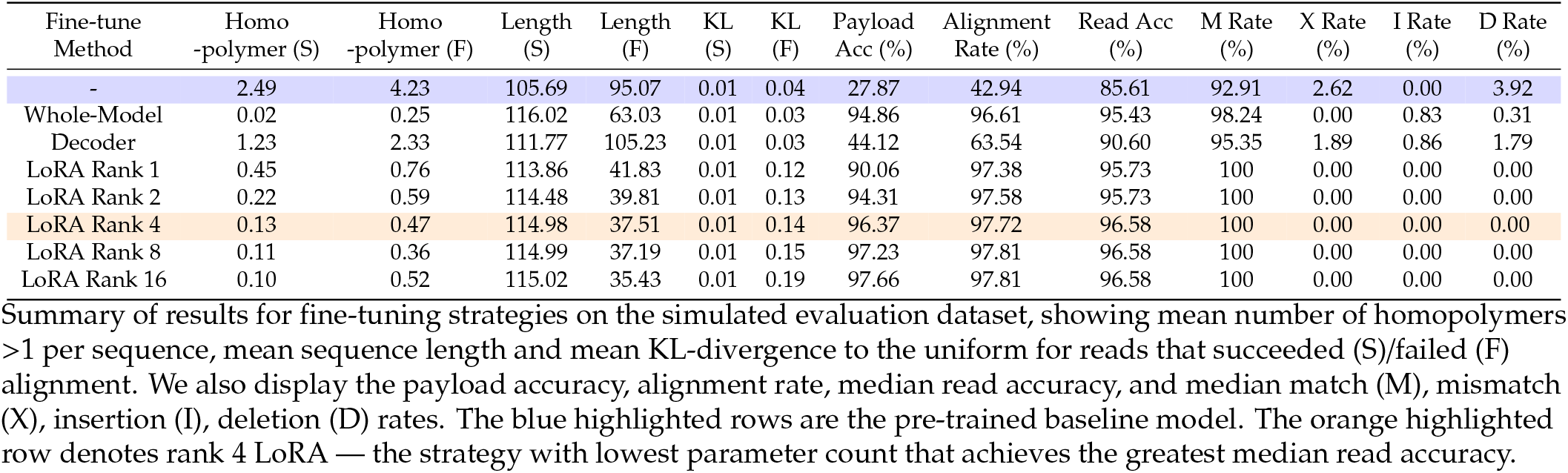
Summary of results for fine-tuning strategies on the simulated evaluation dataset.

**Table 2.**
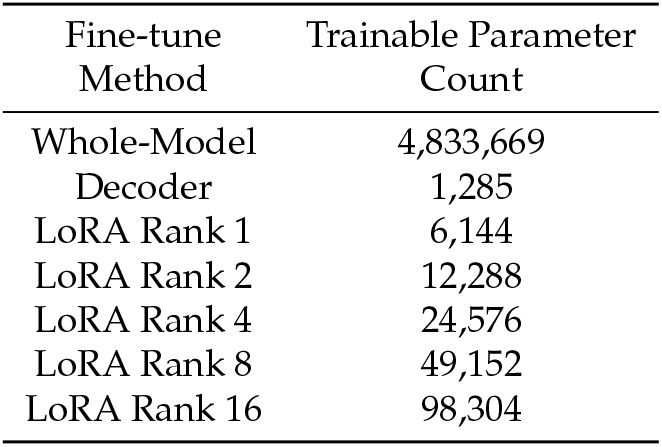
Trainable parameter count of each fine-tuning method.

All LoRA models achieved a median match rate of 100%, even LoRA rank 1, which has the same order of trainable parameters as decoder fine-tuning and only 0.13% of the total model parameter count. This means that half of all alignments produced by Mappy contained no IDS (insertion, deletion or substitution) errors. Increasing rank further (>1) yielded only small improvements of <1% to alignment rate and median read accuracy. However, payload accuracy continued to improve despite this, with LoRA rank 16 achieving the greatest payload accuracy across all files of up to 98%. This highlights that the transformer encoder layer weights have a significantly greater impact on model performance than the linear decoder.

The constraint-related measurements followed a consistent trend. Across all fine-tuning models, the average number of homopolymers (exceeding length 1) and average KL-divergence decreased towards 0, and the average sequence length moved towards 117nt. This shows that fine-tuned models are capable of implicitly learning the constraints present in the data-encoding DNA. As expected, the metrics for sequences that failed alignment were considerably worse, since sequences that do not adhere to the constraints are more likely to have errors. Remarkably, whole-model fine-tuning seems to have learned the constraints better than the LoRA models, with a lower average homopolymer length, sequence length and KL-divergence for successfully aligned sequences, despite a lower alignment rate and read accuracy.

Due to the lack of publicly available datasets for data-encoding nanopore signal data, we also thought it important to further investigate how LoRA performance varied with the amount of training data. We showed (S2 Appendix) that raw signal data corresponding to approximately 1,900 sequences (around 9.5KB of data) is required for LoRA fine-tuning to achieve within approximately 1% of the performance obtained when trained on 933,902 sequences (4.7MB of data). This means that in practice, very little data would need to be synthesised for fine-tuning a basecaller on data-encoding DNA.

It should however be noted that the use of simulated nanopore signal data may not fully capture the noise characteristics and variability of real sequencing experiments. As such, the observed accuracy improvements, may be somewhat inflated compared to performance on actual wet-lab data.

#### Profiling

Portability is an important aspect of nanopore sequencing [7], hence it’s important to consider CPU and GPU utilisation of each fine-tuning method. We profiled all fine-tuning strategies using PyTorch’s built-in profiler, averaging results over ten training steps. Table 3 shows the profiling statistics for each model (averaged across ten training steps).

**Table 3.**
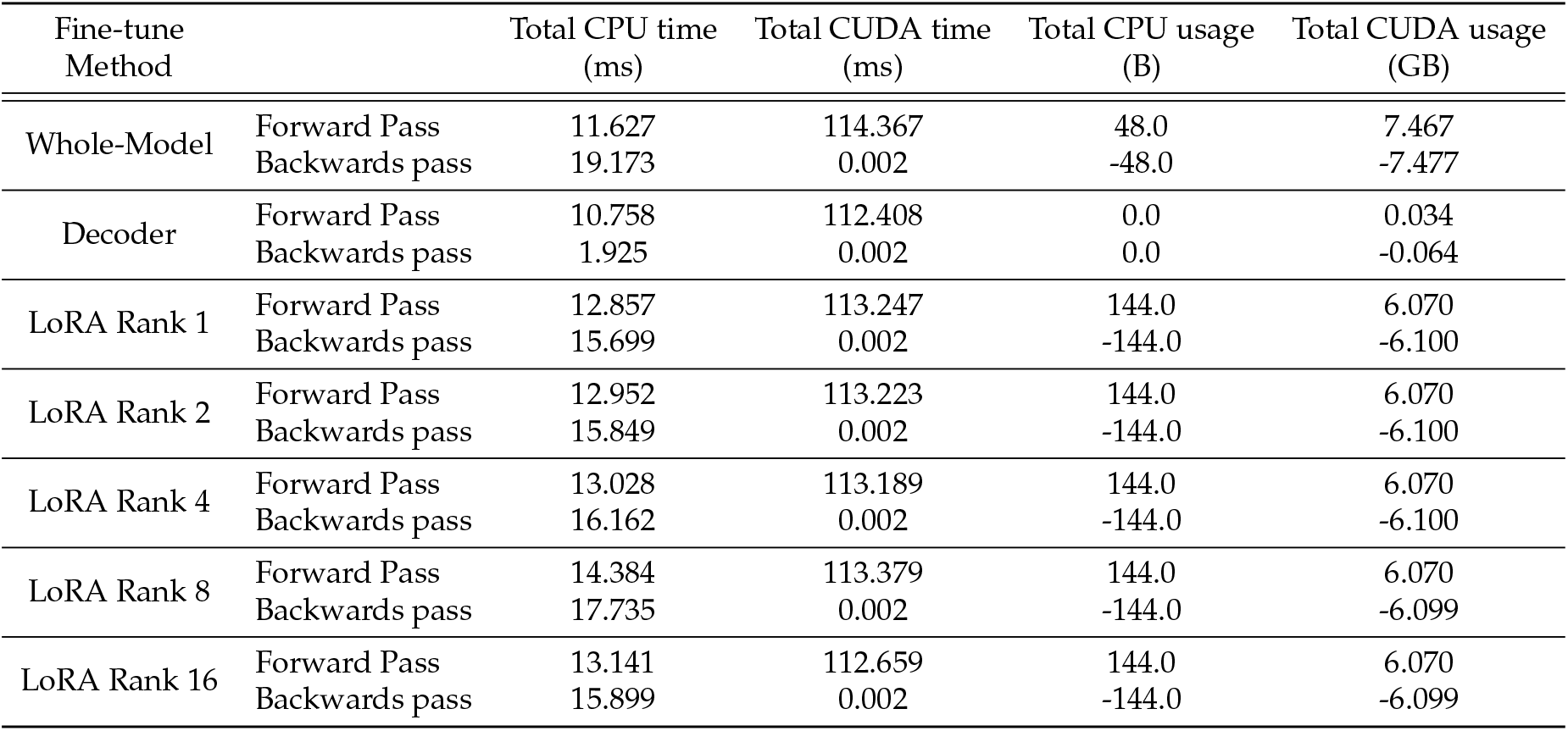
Total CPU time, Total CUDA time, Total CPU usage (RAM), Total CUDA usage (VRAM) broken down by module and averaged over 10 training steps.

Unsurprisingly, decoder fine-tuning exhibited the lowest resource usage across all metrics, utilising only 0.034GB of VRAM, 10.76ms of CPU time and 112.4ms of CUDA time in the forwards pass (and a total of 1.93ms of CPU time in the backwards pass). This is because only 0.03% of the model parameters are trainable and the decoder is the final component of the model — resulting in a small computational graph and minimal parameters that need to be considered for backpropagation.

On the other hand, LoRA had a slightly longer forwards pass than whole-model fine-tuning, which can be attributed to the additional matrix operations required. However, it benefitted from a (up to 1.2 times) faster backwards pass due to reduced parameter count and smaller computational graph. LoRA also consumed approximately 1.5GB (nearly 20%) less VRAM compared to whole-model fine-tuning, which enabled us to fine-tune the model on a laptop’s Nvidia RTX 3080 Mobile GPU with 8GB VRAM; which was not possible for whole-model fine-tuning (or pre-training of the benchmark model). In addition, increasing LoRA rank did not significantly impact CPU or CUDA utilisation.

### Beam Search Evaluation

#### Homopoylmer constraint

Table 4 summarises the performance of our benchmark model on our simulated evaluation dataset for different penalty and max homopolymer length values. Note that a penalty of 1 is equivalent to discarding beams with homopolymers.

**Table 4.**
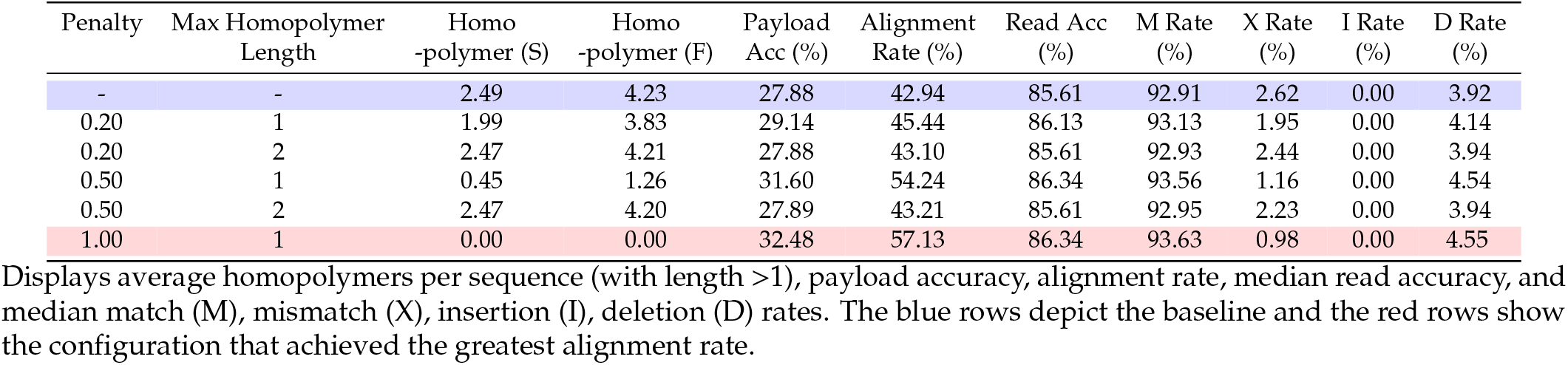
Summary of results with beam search modifications to penalise extensions exceeding a maximum homopolymer length.

As the results show, discarding any hypothesis extensions violating the maximum homopolymer length significantly increased the alignment rate by 13-22%, median read accuracy by 0.85% and payload accuracy by 3.63-7.47%. Another important observation is that using a maximum homopolymer length of 2 when penalising extensions had little effect on the average number of homopolymers per sequence for both successful and failed alignments. This shows that the basecaller very rarely predicts homopolymers with length greater than 2. This, combined with a significantly lower mean number of homopolymers per sequence when using a maximum homopolymer length of 1, implies that the majority of homopolymers predicted by the basecaller were actually of length 2 — further explaining why this value of maximum homopolymer length performed poorly compared to a value of 1.

#### Length constraint

Table 5 shows the performance of the model when penalising by length for different values of λ. Table 6 shows performance of the model when discarding beams that differ from the desired length per number of time steps for different thresholds θ.

The first strategy, which penalised extensions, performed better overall, achieving up to a 3% increase in alignment rate, whereas discarding extensions actually decreased it by up to 0.6%. Despite this, the latter strategy had a more pronounced effect on the average length of decoded nucleotide sequences, with an increase of approximately 5nt, whereas penalising extensions only increased the average length of sequences by 0.85-2nt (with the best configuration). This indicates that the probability distributions predicted by the benchmark basecaller limits the effectiveness of these modifications, as although our second strategy generated sequences that adhered more closely to the 117nt length constraint, the alignment rate and payload accuracy gains were much lower than those obtained from the first strategy.

This is reinforced by examining the IDS (insertion, deletion and substitution) error rates. Penalising extensions had little effect on the median mismatch rate, but did decrease the median deletion rate. This means that more bases in the reference sequence were aligned to the predicted sequence. However, using a penalty coefficient greater than 15 resulted in insertion errors, reducing the overall median match rate despite continuing to reduce the number of deletions. When discarding extensions, minimal changes to the median match rate or IDS rates were observed despite the more noticeable increase in mean predicted sequence length. This implies that Mappy excluded the extra predicted bases from alignment, because they were too erroneous, thus explaining why this strategy actually decreased alignment rate.

**Table 5.**
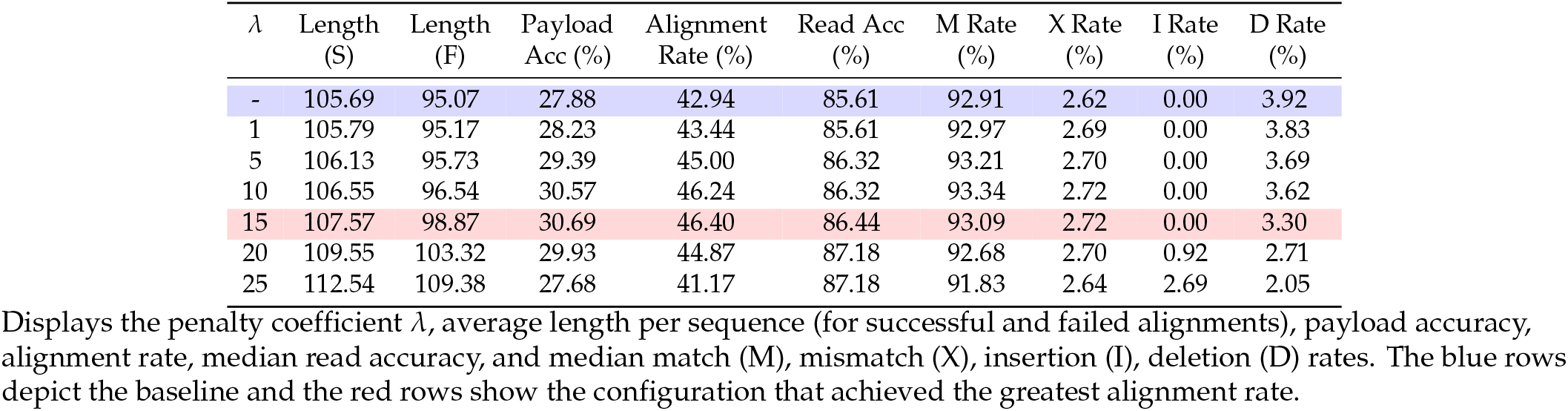
Summary of results with beam search modifications to penalise extensions by their normalised length.

**Table 6.**
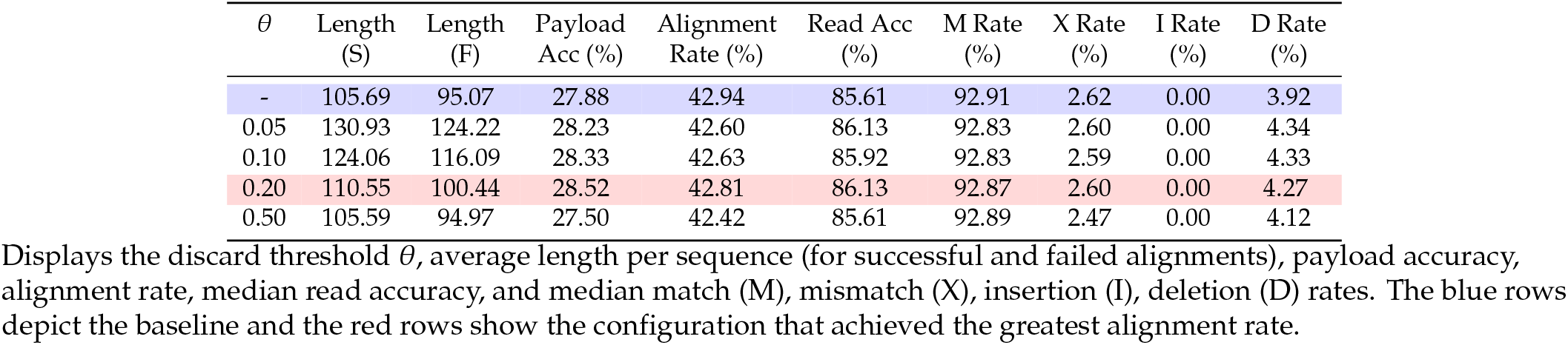
Summary of results with beam search modifications to discard extensions that are too short at a time step.

Overall, leveraging the length constraint is far less effective than leveraging the maximum homopolymer constraint. Although both methods are capable of small median read accuracy increases (<1%), our attempts at leveraging the length constraint only improved payload accuracy by <3%, compared to the 3.63-7.47% increase observed when leveraging the maximum homopolymer constraint.

#### Base distribution constraint

Table 7 shows the performance of the model when penalising by KL-divergence for different values of λ. Table 8 shows the performance of the model when only considering under-represented beam extensions for different values of θ.

Implementing a KL-divergence penalty into the scoring function achieved slight improvements to the alignment rate and payload accuracy of approximately 1%. However, for higher penalty coefficients, the median read accuracy decreased and at λ = 20, the alignment rate was so low that very few, if any decoded sequences were successfully aligned, despite having a low KL-divergence. This is due to the algorithm overly favouring sequences with low KL-divergence, resulting only in decoded sequences that were permutations of ‘ACGT’ and thus no successful alignments.

**Table 7.**
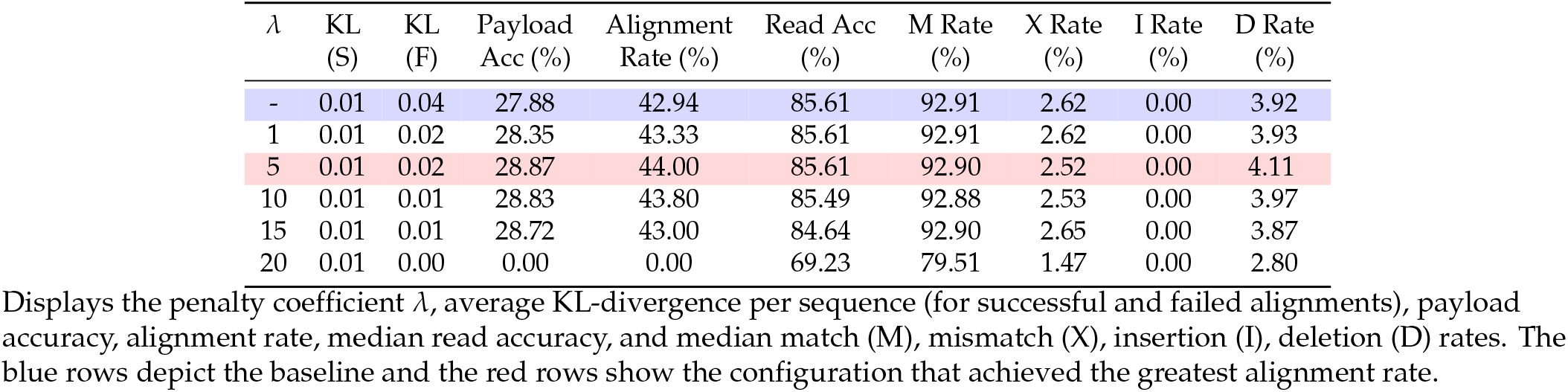
Summary of results with beam search modifications to penalise extensions by the KL-divergence of the base distribution to the uniform.

**Table 8.**
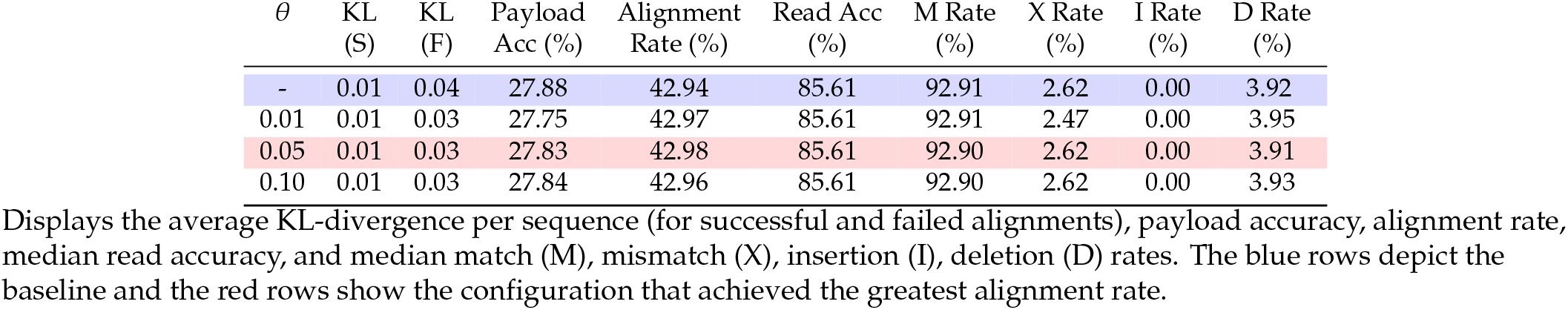
Summary of results with beam search modifications to discard extensions exceeding a KL-divergence threshold of θ and entropy threshold of 0.8 · log 4.

On the other hand, discarding extensions performed worse, with alignment rate improvements of merely <0.1% and often worse payload accuracies. Furthermore, the average KL-divergence of the base distribution of sequences to the uniform changed minimally. This means very few extensions were actually discarded, due to not exceeding the entropy threshold.

Overall, it has been shown that it is possible to leverage the base distribution constraint using a KL-divergence penalty to achieve alignment rate improvements. However, its effectiveness is limited since these methods rely on the correctness of the basecaller output. In addition, the KL-divergence of a sequence to the uniform distribution depends on all previous predicted nucleotides. This means that incorrect predictions have a knock-on effect, impacting future predictions and potentially resulting in more incorrect nucleotides being selected purely to reduce the KL-divergence.

#### Combining Constraints

We investigated leveraging all three constraints by combining the best performing strategy for each one:

1. Discarding any extension if it would contain a homopolymer of length >1
2. Applying a normalised length penalty
3. Applying a KL-divergence penalty

Table 9 displays the performance for varying length λ_len_ and KL λ_KL_ penalty coefficients.

**Table 9.**
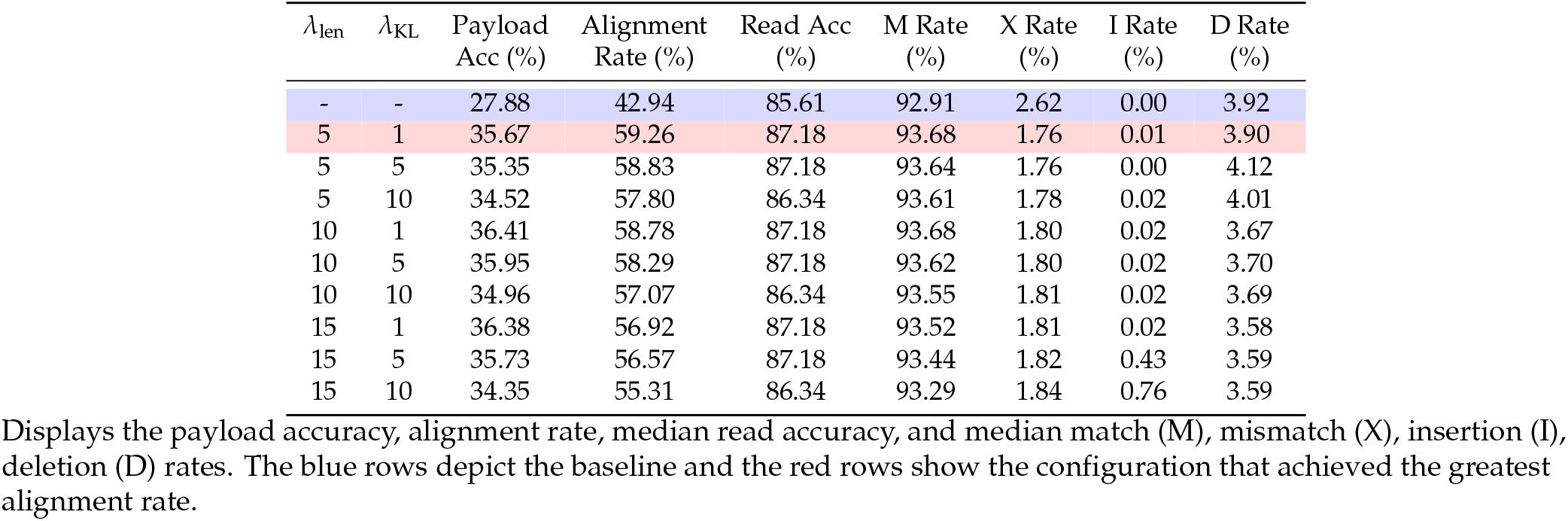
Summary of results when discarding homopolymers of length >1, and varying the normalised length λ_len_ and KL penalty coefficients λ_KL_.

The best performing configuration across all files in terms of alignment rate was λ_len_ = 5 and λ_KL_ = 1, increasing alignment rate by approximately 15-22%, and increasing median read accuracy by up to 1.7%, more than double the 0.75% increase observed by Sneddon et al. with their language-model informed beam search [21]. While this configuration did not necessarily yield the greatest payload accuracy, it improved upon the baseline by 6.63-11.3%. In addition, it outperformed the best individually applied constraint (discarding illegal homopolymers) in all metrics, implying that leveraging multiple constraints simultaneously has a synergistic effect — improving performance gains beyond what any single strategy could achieve.

Furthermore, we explored the impact of the best parameter configuration on the previously fine-tuned model. The results (S4 Table) show negligible improvements to alignment rate and payload accuracy (<1%) for all strategies other than decoder fine-tuning, which experienced gains of >5% and 1.58-2.79% respectively. The modifications also marginally improved read accuracy in some cases, most notably, increasing the median read accuracy of LoRA rank 2 from 95.73% to 96.58% — the same read accuracy achieved by LoRA rank 4. As these improvements are only slight, it reinforces that fine-tuning with LoRA allows the basecaller to effectively learn the constraints present in data-encoding DNA.

## Conclusions

This work explored the extent to which constraints imposed on data-encoding DNA can be incorporated into basecalling to improve accuracy in a DNA storage context. By investigating two methods that can be applied to the majority of available, pretrained basecallers - parameter-efficient fine-tuning and constraint-aware decoding - we evaluated how structural features of data-encoding DNA, such as limited homopolymers and balanced base composition, can be exploited to increase both read accuracy and overall data recovery.

Fine-tuning with Low-Rank Adaptation (LoRA) emerged as a particularly effective strategy. Updating just a small fraction of the model parameters (as little as 0.25%) improved median read accuracy from 85.8% to over 95%, and increased data recovery rates to more than 94%, approximately 65% more than the pre-trained model. A higher-rank LoRA configuration of 16 achieved over 97% data recovery without any external error correction, all while reducing memory usage relative to whole-model fine-tuning. Notably, these gains were achieved with as few as 1,900 synthetic DNA strands (∼9.5 kB), indicating that very little data would need to be synthesised to fine-tune a basecaller on data-encoding DNA. This result aligns with earlier work by Pagès-Gallego et al. [17], which similarly emphasized the practicality of fine-tuning on limited samples.

In parallel, we explored a constraint-aware decoding approach that penalises violations of homopolymer limits, sequence length, and base composition balance. Without requiring any retraining, this method increased alignment rates by up to 22% and improved read accuracy by 1.7%. These improvements substantially exceed those reported in earlier decoder-based interventions, such as the language-informed beam search by Sneddon et al. for RNA sequencing, which achieved only a 0.75% gain [21]. Furthermore, the modifications to the algorithm do not impact the overall time complexity and require no re-training of the model - this makes it attractive in scenarios where there is no training data or hardware is a limiting factor.

These results provide strong evidence that the inherent structure of data-encoding DNA enables more effective optimisation of basecalling than traditional biological sequencing. By improving basecalling accuracy, these methods also reduce the need for heavy error-correction codes such as HEDGES [30], thereby increasing information density and lowering the computational cost—both critical factors for building scalable and efficient DNA data storage technologies.

## Future work

Future work should validate these findings on real, synthesised data-encoding DNA to assess robustness in practical scenarios. Extending the evaluation to other coding schemes and basecaller architectures would also help assess generalisability. Moreover, exploring alternative parameter-efficient fine-tuning methods, such as adapter tuning [31], prefix-tuning [32] and BitFit [33], could reveal further trade-offs in performance, speed, and memory use. Lastly, given the structured nature of synthetic DNA, there is strong potential for developing specialised, lightweight basecallers or even universal models trained on multiple coding schemes, enabling broad applicability with minimal fine-tuning requirements.

## Supporting information

**S1 Table Details on the test set encoded with the Goldman code [9]**.

**S2 Appendix Leveraging Constraints Through Fine-Tuning - Data regimes**

**S3 Fig. Whole-Model Fine-Tuning Training Curves**

**S4 Table Different fine-tuning methods with and without using the best configuration of combined beam search modifications**.

