## Supplementary material for "Basecalling for DNA Storage": S1 Table

**S1 Table Details on the test set encoded with the Goldman code [1].**

| File | General Information | Number of Sequences |
|---|---|---|
| EBI.jp2 | a photo of the European Bioinformatics Institute in JPEG 2000 format | 37422 |
| MLK_excerpt_VBR_45-85.mp3 | a 26 second excerpt from Martin Luther King's 'I have a dream' speech | 34163 |
| View_huff3.cd | the Huffman code used for the encoding | 3218 |
| watsoncrick.pdf | Watson and Crick's paper on the structure of DNA | 56910 |
| wssnt10.txt | a text file containing all 154 of Shakespeare's sonnets | 21649 |

Includes general file information and number of DNA sequences used to encode the data.
