## Supplementary material for "Basecalling for DNA Storage": S2 Appendix

### S2 Appendix Leveraging Constraints Through Fine-Tuning - Data regimes

Due to the lack of publicly available datasets for data-encoding nanopore signal data, we also thought it important to investigate how LoRA performance varied with the amount of training data. We chose to use rank 4 LoRA, as it was the strategy with lowest trainable parameters that achieved the greatest median read accuracy. Fig 1 depicts the alignment rate, median read accuracy and payload accuracy of rank 4 LoRA (trained for 45,000 training steps, with a batch size of 64) on various data regimes.

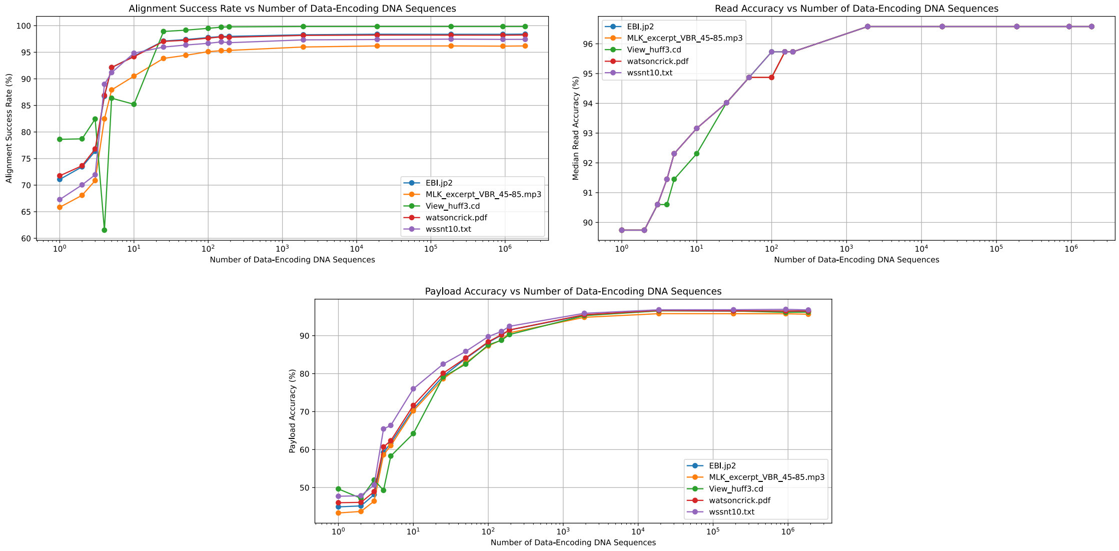


**Fig 1.** The alignment rate, median read, and payload accuracy against number of sequences that the benchmark model was fine-tuned on using rank 4 LoRA

Both alignment rate and read accuracy increased with number of DNA sequences in the dataset, plateauing at around 1,914 sequences, at which point the payload accuracy is within 1% of its peak value. This is a surprising result, as it corresponds to only around 9.5KB of data, whereas our initial fine-tuning experiments were conducted on roughly 4.7MB of data. Furthermore, training the model on only a single data-encoding DNA sequence exceeded the baseline alignment rate by approximately 25%, median read accuracy by >4% and payload accuracy by approximately 16%. This shows that knowledge learned by the model in its pre-training task on bacterial data is heavily transferable to data-encoding DNA. In addition, the small amount of training data required to achieve maximal performance reinforces our hypothesis that basecalling constrained, data-encoding DNA is a fundamentally simpler task than basecalling bacterial DNA.

For completeness, we fine-tuned all tested LoRA ranks on 1,914 DNA sequences. All models achieved the same median read accuracy and within 1% of the alignment rate and payload accuracy as those obtained when fine-tuning on 933,902 sequences.
