## Supplementary material for "Basecalling for DNA Storage": S3 Figure

### S3 Fig. Whole-Model Fine-Tuning Training Curves

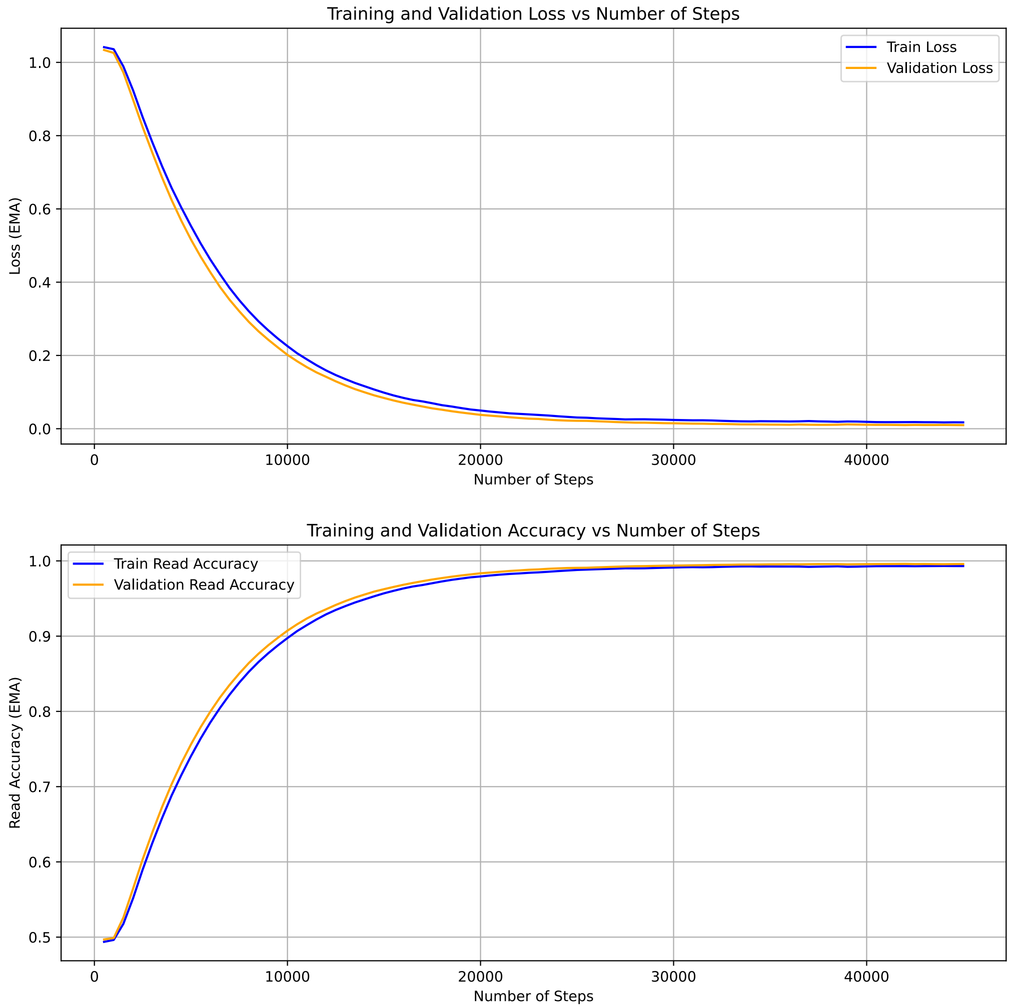


**Fig 1.** The exponential moving average training and validation loss and accuracy for whole-model fine-tuning
