## Supplementary material for "Basecalling for DNA Storage": S4 Table

**S4 Table Different fine-tuning methods with and without using the best configuration of combined beam search modifications.**

| Fine-tune Method | Payload Acc (%) | Alignment Rate (%) | Read Acc (%) | M Rate (%) | X Rate (%) | I Rate (%) | D Rate (%) |
|---|---|---|---|---|---|---|---|
| Whole-Model | 94.84 | 96.61 | 95.43 | 98.24 | 0.00 | 0.83 | 0.31 |
| | 94.91 | 96.61 | 95.75 | 98.24 | 0.00 | 0.84 | 0.00 |
| Decoder | 44.13 | 63.54 | 90.60 | 95.35 | 1.89 | 0.86 | 1.79 |
| | 46.55 | 69.20 | 90.62 | 95.52 | 1.80 | 0.88 | 1.66 |
| LoRA rank 1 | 90.09 | 97.38 | 95.73 | 100.00 | 0.00 | 0.00 | 0.00 |
| | 91.76 | 97.48 | 95.73 | 100.00 | 0.00 | 0.00 | 0.00 |
| LoRA rank 2 | 94.30 | 97.58 | 95.73 | 100.00 | 0.00 | 0.00 | 0.00 |
| | 95.02 | 97.62 | 96.58 | 100.00 | 0.00 | 0.00 | 0.00 |
| LoRA rank 4 | 96.38 | 97.72 | 96.58 | 100.00 | 0.00 | 0.00 | 0.00 |
| | 96.71 | 97.75 | 96.58 | 100.00 | 0.00 | 0.00 | 0.00 |
| LoRA rank 8 | 97.22 | 97.81 | 96.58 | 100.00 | 0.00 | 0.00 | 0.00 |
| | 97.43 | 97.83 | 96.58 | 100.00 | 0.00 | 0.00 | 0.00 |
| LoRA rank 16 | 97.68 | 97.81 | 96.58 | 100.00 | 0.00 | 0.00 | 0.00 |
| | 97.91 | 97.83 | 96.58 | 100.00 | 0.00 | 0.00 | 0.00 |

The blue highlighted rows are the baseline without any modifications. Displays payload accuracy, alignment rate, median read accuracy and median match (M)/mismatch (X)/insertion (I)/deletion (D) rates
